# Spine-Prints: Transposing Brain Fingerprints to the Spinal Cord

**DOI:** 10.1101/2025.05.30.656545

**Authors:** Ilaria Ricchi, Andrea Santoro, Nawal Kinany, Caroline Landelle, Ali Khatibi, Shahabeddin Vahdat, Julien Doyon, Robert L. Barry, Dimitri Van De Ville

## Abstract

Functional connectivity (FC) patterns in the human brain form a reproducible, individual-specific “fingerprint” that allows reliable identification of the same participant across scans acquired over different sessions. While brain fingerprinting is robust across healthy individuals and neuroimaging modalities, little is known about whether the fingerprinting principle extends beyond the brain. Here, we used multiple functional magnetic resonance imaging (fMRI) datasets acquired at different sites to examine whether a fingerprint can be revealed from FCs of the cervical region of the human spinal cord. Our results demonstrate that the functional organisation of this region also exhibits individual-specific properties, suggesting the potential existence of a spine-print within the same acquisition session. Although the spine-print scores are not directly comparable to those observed in the brain, this discrepancy may in part reflect the intrinsic limitations of imaging this region with fMRI, where where the signals are more susceptible to noise and effective resolution relative to structure size, and tSNR are markedly lower than in the brain.This study provides the first evidence of a spinal cord connectivity fingerprint, underscoring the importance of considering a more comprehensive view of the entire central nervous system. Eventually, these spine-specific signatures could contribute to identifying individualized biomarkers of neuronal connectivity, with potential clinical applications in neurology and neurosurgery.

## 1. Introduction

Functional magnetic resonance imaging (fMRI) provides a non-invasive tool for capturing the spatiotemporal features of brain activity across individuals (Dubois et al., 2024; McGonigle et al., 2000). For resting state, pairwise correlations between regionally-averaged time series define a functional connectivity (FC) profile that characterizes statistical dependencies between activity in different brain regions (Fornito et al., 2016; Sporns et al., 2005). While most of the early studies in neuroimaging have analyzed FC patterns at the group level to identify robust population trends (Margulies et al., 2016; Snyder & Raichle, 2012), such approaches might overlook meaningful inter-individual differences. In the human brain, FC profiles have been shown to act as distinctive neural “fingerprints”. The pivotal study of Finn et al., 2015 demonstrated that each individual’s FC pattern remains consistent across multiple sessions and conditions, allowing to reach up to 95% identification accuracy across 126 subjects at rest. In this context, the frontoparietal network emerged as being particularly distinctive, hence enabling accurate identification of individualized FC patterns, underscoring their potential for personalized neuroimaging applications. Over the past decade, a growing body of work has shown that the accuracy of brain fingerprinting depends on multiple factors, including scan length, the inter-scan interval (Amico & Goñi, 2018; Horien et al., 2019) and the presence of different clinical (Sorrentino et al., 2023; Stampacchia et al., 2024, 2025) or cognitive conditions (Luppi et al., 2022, 2024; Tolle et al., 2024). Methodological advances have also improved performance. For instance, Amico & Goñi, 2018 examined approaches to enhance the distinctiveness of individual functional connectomes using dimensionality reduction based on principal component analysis (PCA). They quantified how consistently an individual’s FC profile could be recognized across different sessions, achieving near-perfect classification when only the most informative connections were retained.

Similarly, Li et al., 2021 investigated optimal strategies for feature extraction in brain fingerprinting, by selecting the most relevant edges rather than the whole brain connections. In terms of the dynamic and temporal features of brain fingerprints, Van De Ville et al., 2021 showed that identification is possible even at short time scales (*i.e.* in less than 30 seconds of fMRI activity), but that the strength fluctuates significantly over time. In a more recent study, Ghaffari et al., 2025 demonstrated that the subcortical-cerebellum network had a substantial contribution in the static case, while playing a less prominent role in any of the dynamic fingerprints. Other studies have also investigated an individualized approach using Connectome Fingerprint (Tobyne et al., 2018), which is more focused on the prediction of individualized task activation. Building on this work, Tripathi & Somers, 2023 found that using a subject’s own FC generally resulted in higher prediction accuracy for their task activations compared to using connectivity from other individuals. In particular, FC patterns between the cerebral cortex and cerebellum carried crucial information for these predictions. A minor involvement of the cerebellum for the fingerprint accuracy was also found (Mantwill et al., 2022). Beyond fMRI, brain fingerprinting extends across various neuroimaging modalities, including electroencephalography (EEG) (Demuru & Fraschini, 2020; Fraschini et al., 2016; W. Kong et al., 2019), functional near-infrared spectroscopy (fNIRS) (de Souza Rodrigues et al., 2019), and magnetoencephalography (MEG) (da Silva Castanheira et al., 2021; Sareen et al., 2021), highlighting the robustness of individual-specific connectivity patterns across different measurement techniques.

While brain functional connectivity has been widely studied, the same cannot be said for the spinal cord, despite its pivotal role in sensory and motor functions (for reviews Landelle et al., 2021; Kinany, 2022). However, the spinal cord is more than a pathway for relaying sensory and motor signals between the brain and the effectors. A large body of literature demonstrates that the spinal cord is also involved in a range of integrative and plastic processes, including learning (Grau, 2014; Khatibi et al., 2022; Vahdat et al., 2015; Wolpaw, 2012), pain modulation, and modulation of descending signal from supraspinal structures (Ossipov et al., 2014; Martucci & Mackey, 2018; Nim et al., 2021). Furthermore, a series of pioneering studies (Khatibi et al., 2022; Kinany et al., 2023; Vahdat et al., 2015) investigating motor sequence learning with simultaneous human brain and spinal cord fMRI have provided *in vivo* evidence supporting learning-related plasticity within the spinal cord. Complementing this, more recent findings suggest that the human spinal cord is also involved in predictive processing, with prior knowledge influencing spinal responses to sensory stimuli as early as 13–16 milliseconds after stimulation (Stenner et al., 2025). These insights are further supported by neurophysiological evidence pointing to a complexity in spinal function beyond its classical sensorimotor role, including contributions to affective dimensions. For example, observing others in pain can evoke spinal responses similar to those triggered by first hand pain (Tinnermann, 2021). Moreover, local interneuronal circuits appear to support task learning, and computational models suggest that these circuits may self-organize through Hebbian learning (Enander et al., 2022). Altogether, these processes may give rise to nuanced, individual-specific connectivity patterns in the spinal cord. Supporting this idea, test-retest experiments have shown that functional connectivity profiles remain stable across sessions, suggesting consistent and potentially unique signatures at the individual level—a prerequisite to probe the existence of a “spinal cord fingerprint”, or “spine-print” (Barry et al., 2016; Kaptan et al., 2023; Kowalczyk et al., 2024). Importantly, these prior studies primarily addressed intrasubject reliability, whereas fingerprinting shifts the focus toward intersubject identifiability, assessing whether an individual can be distinguished from others based on their connectivity profile. In this sense, fingerprinting builds upon reliability but extends it toward inter-individual distinctiveness. Furthermore, reliability typically evaluates the stability of individual connections in a univariate fashion, whereas fingerprinting captures multivariate patterns across the whole connectome, yielding a richer and more distinctive individual signature.Inspired by brain fingerprinting and the spinal-cord reliability studies, herein we provide the first direct evidence supporting the existence of a spinal cord fingerprint, for which we coin the term “spine-print”. We investigate whether individual FC patterns could be reliably identified in the cervical spinal cord, akin to those observed in the brain. To test this, we computed identifiability scores from cervical spinal cord FC profiles using three independent datasets, one of which included simultaneous brain and spinal cord acquisitions. Following Amico & Goñi, 2018, we implemented a pipeline to extract identifiability metrics and assess the reliability of specific spinal connections, and successfully confirmed the existence of spine-prints including which functional connections were most distinctive across individuals. In addition to assessing spinal cord identifiability in isolation, we established the individual fingerprints of the brain, the spinal cord, and their interaction. These findings contribute to a growing body of work suggesting that the spinal cord, far from being a mere relay, may play a dynamic and individual-specific role in the human central nervous system.

## 2. Methods

### 2.1. Datasets

This study examines three distinct datasets: *Dataset 1* (partially published data from Vanderbilt University), which originally comprised 20 healthy volunteers (10 female, mean age 32 ± 11 years). However, two female participants were excluded—one due to missing physiological recordings and the other because of a higher slice placement along the spinal cord, which complicated normalization to the template and reduced consistency in spinal level overlap across participants; *Dataset 2* from the Max Planck Institute in Leipzig, Germany that consists of 43 participants selected from a pool of 48 subjects in the publicly available OpenNeuro data (Kaptan et al., 2023): five participants were excluded (four due to missing physiological data and one due to a different number of acquired volumes); Dataset 3 (unpublished data with similar acquisition protocol to Landelle et al., 2024) includes 15 participants (7 female, 30±6 years old).

Participants in *Dataset 1* provided written informed consent under a protocol approved by the Vanderbilt University Institutional Review Board. These data were then analyzed under a protocol approved by the Mass General Brigham Institutional Review Board. *Dataset 2* was already published (Kaptan et al., 2023), having therefore ethical approval. *Dataset 3* study was approved by the ethic committee of the Institut universitaire de gériatrie de Montréal (#CERVN 17-18-20), which follows the policies of the Canadian Tri-Council Research Ethics Policy Statement and the principles expressed in the Declaration of Helsinki.

### 2.2. Data acquisition

#### Dataset 1

These data were acquired by R. Barry while at the Vanderbilt University Institute of Imaging Science. Experiments were performed on a Philips Achieva 3T scanner (Best, The Netherlands) with a dual-channel transmit body coil and the vendor’s 16-channel neurovascular coil (6 head, 4 neck, and 6 upper chest channels). Each scanning session began with a sagittal localizer to identify the general anatomy and location of the C3/C4 intervertebral disc. The imaging stack was then centered at this level, ensuring that all slices were perpendicular to the cord. The imaging stack covered vertebral levels C2-C5, roughly corresponding to the spinal nerve root levels C3-C6. High-resolution axial anatomical images were acquired using an averaged multi-echo gradient echo (mFFE) T2*-weighted sequence with the following parameters: field of view (FOV) = 150 × 150 mm^2^, acquired voxel size = 0.65 × 0.65 × 5 mm^3^, interpolated voxel size = 0.29 × 0.29 × 5 mm^3^, 12 slices, first TE = 7.20 ms, 4 additional echoes where ΔTE = 8.83 ms (5 echoes in total), repetition time (TR) = 700 ms, flip angle = 28°, sensitivity encoding (SENSE) = 2.0 (left-right), and number of acquisitions averaged = 2. Total acquisition time for the anatomical scan was 5 mins and 26 s. A saturation band was positioned anterior to the spinal cord to suppress signal from the mouth and throat. These anatomical data were presented in a previous publication measuring T2* in spinal cord gray matter (Barry & Smith, 2019). Resting state fMRI data were acquired with identical slice placement using a 3D multi-shot gradient-echo sequence: volume acquisition time (VAT) = 2080 ms, TR/TE= 34/8 ms, flip angle = 8°, voxel size = 1 × 1 × 5 mm^3^, 12 slices. The two separate 10-min (288 volumes each) were separated by a ∼2-min period to give the subject a chance to rest. Respiratory and cardiac cycles throughout both runs were externally monitored and continuously recorded using a respiratory bellows and a pulse oximeter. These fMRI data were previously analyzed and presented in a conference abstract (Rangaprakash & Barry, 2021).

#### Dataset 2

The second dataset, described in detail in Kaptan et al., 2023, was acquired using a 3T Siemens Prisma MRI system equipped with a whole-body RF coil, a 64-channel head-and-neck coil, and a 32-channel spine-array coil. Participants were instructed to remain still, breathe normally, and avoid excessive swallowing. The dataset is part of a larger methodological project, focusing on two functional MRI acquisitions and one structural acquisition.

Functional MRI runs included 250 single-shot gradient-echo EPI volumes (VAT = TR × number of shots (1) = 2312 ms, 9.63 min of duration), covering the spinal cord from C2 to T1 with 24 slices (5 mm thickness) and a cross-section resolution of 1.0 × 1.0 mm². The acquisitions used z-shimming to counteract signal loss due to magnetic field inhomogeneities, with two runs differing in manual vs. automatic z-shim selection (Kaptan et al., 2022). The two runs were separated by a maximum of ∼10 min.

A high-resolution T2-weighted 3D sagittal SPACE sequence (0.8 × 0.8 × 0.8 mm³) was acquired for registration. Peripheral physiological signals (respiration via a breathing belt and cardiac activity via ECG electrodes) were also recorded for physiological noise modeling.

#### Dataset 3

The last dataset was acquired by J. Doyon’s group. It is characterized by 2 functional runs back-to-back, also intervalled by a few minutes following similar acquisition parameters as Landelle et al. 2024. Data were acquired on a 3T MRI Scanner (Magnetom-Prisma, Siemens, Erlangen, Germany) with a 64 channel head and neck coil. The blood-oxygen-level-dependant (BOLD) images were acquired using a gradient-echo echo-planar imaging (EPI) sequence with the following parameters: VAT = TR = 1550 ms, TE = 23 ms, in-plane voxel resolution = 1.6 × 1.6 mm^2^; slice thickness = 4 mm, FOV = 120 mm x 120 mm, flip angle = 70°; iPAT acceleration factor PE = 2, iPAT acceleration factor slice = 3. A total of 69 axial slices were acquired from the top head to C7/T1 vertebrae. Each resting-state run lasted 10 minutes and 35 seconds (230 volumes). Physiological recordings were acquired using a pulse sensor and a respiration belt (Siemens Physiology Monitoring Unit).

Brain/spinal cord structural images were acquired using a high-resolution T1-weighted anatomical image in the sagittal direction (MPRAGE sequence: TR/TE = 2300/3.3 ms, voxel size = 1.3 × 1.3 × 1.3 mm^3^, FOV = 365 x 375 mm^2^, flip angle = 9°). In total, 288 slices were acquired, covering the top of the head to the upper thoracic regions T2-weighted images were acquired, spanning from the top of the cerebellum to the upper thoracic region (approximately at T1), thereby encompassing the entirety of the cervical spinal cord. These images were collected in the transverse orientation with the following parameters: TR = 33 ms; TE = 14 ms; FOV = 211 mm × 249 mm; flip angle = 5°; in-plane voxel resolution = 0.35 mm × 0.35 mm, slice thickness = 2 mm.

### 2.3. Data preprocessing

We applied an in-house preprocessing pipeline for the first two spinal cord datasets publicly available on github (https://github.com/MIPLabCH/SC-Preprocessing.git), while the third brain/spinal cord dataset was preprocessed using (Landelle et al., 2024) preprocessing pipeline. The two pipelines include comparable preprocessing steps for the spinal cord; see (Landelle et al., 2024; Ricchi, Kinany, et al., 2024) for reference, no spatial smoothing was applied to minimize partial volume effects, which is standard practice in spinal cord studies given the small ROI dimensions (Barry et al., 2014, 2018; Eippert et al., 2017).

#### 2.3.1. Preprocessing of Datasets 1 & 2

The spinal cord functional and structural images were pre-processed using Python (version 3.9.19), with the nilearn library (version 0.9.1) falling under the umbrella of scikit-learn (version 0.24.2), FMRIB Software Library (FSL; version 5.0), and Spinal Cord Toolbox (SCT; version 5.3.0; (De Leener et al., 2017)). The following preprocessing steps were performed: i) slice-timing correction (FSL, *‘slicetimer’*), ii) motion correction using slice-wise realignment and spline interpolation (with SCT, ‘*sct_fmri_moco’*), iii) segmentation of functional and structural images (with ‘*sct_deepseg’*, followed by manual correction), iv) time series denoising (see details in the next paragraph), v) registration of functional images to anatomical images, and finally vi) coregistration of functional images to anatomical images, and then, to the PAM50 template (with SCT, ‘*sct_register’* multimodal and ‘*sct_register_to_template’* with nearest neighbor interpolation). Additionally, we assessed the data quality of the three datasets by obtaining voxelwise tSNR values with the SCT’s function *‘sct_fmri_compute_tsnr’* (De Leener et al., 2017), which computes each voxel’s temporal mean and divides it by its standard deviation.

#### 2.3.2. Time series denoising of Datasets 1 & 2

The time series denoising follows the same procedure as previous spinal fMRI (Kinany et al., 2024; Landelle et al., 2023, 2024; Ricchi, Kinany, et al., 2024) relying on the *‘clean_img*’ function from the nilearn library, This approach enable the removal of the noise confounds orthogonally to the temporal filter. Specifically, confounds and the band-pass temporal filter (cut-off frequencies: 0.01 Hz and 0.13 Hz) were projected onto the same orthogonal space, following the methodology outlined in Lindquist et al., 2019, instead of being applied sequentially.

Physiological noise correction was performed using a model-based approach inspired by RETROICOR (RETROspective Image CORrection; (Glover, 1999)), which models physiological signals as quasi-periodic and maps their phases onto each image volume using a Fourier series expansion. To implement this, we used FSL’s Physiological Noise Modeling (PNM) tool to generate nuisance regressors from cardiac, respiratory, and cerebrospinal fluid (CSF) signals. Cardiac peaks were identified using the *‘scipy.signal.find_peaks’* function (Virtanen et al., 2020), followed by manual verification to ensure detection accuracy.

We followed established guidelines for physiological noise correction in spinal cord fMRI (Y. Kong et al., 2012). Both cardiac and respiratory regressors were modeled with an order of 4, including the fundamental frequency and the first three harmonics. Their interaction was modeled up to second order, producing a total of 32 regressors on a slice-by-slice basis. Additionally, a CSF regressor was derived by averaging the signal from the 10% most variable CSF voxels, and the estimated motion correction parameters were included as regressors. Each of these regressors was applied independently per slice using the *‘clean_img*’ function to account for the anatomical and physiological specificity of spinal cord data.

#### 2.3.3. Preprocessing and denoising of Dataset 3

As this dataset includes simultaneous brain and spinal cord acquisition, we adapted the previously described approach accordingly. For each participant, all the functional and structural images were pre-processed separately for the brain and the spinal cord using the Spinal Cord Toolbox (SCT; version 5.6), FSL, SPM, and in-house Python and MATLAB programs. First, T1w images were cropped to separate the brain and spinal cord. Spinal cord segmentation was carried out using SCT (sct_propseg) and manually corrected when necessary. The brain structural images were segmented using Cat-12 toolbox (SPM) into gray matter (GM), white matter (WM), and CSF.

The spinal cord volumes of each functional run were averaged, and the centerline of the cord was automatically extracted from the mean image (or drawn manually when needed). A cylindrical coarse mask with a diameter of 15 mm along this centerline was drawn and further used to exclude regions outside the spinal cord from the motion correction procedure, as those regions may move independently from the cord. Motion correction was performed using the first volume as the target image and resulted in the motion corrected time series (‘*sct_fmri_moco’* from SCT). Moco was carried out using slice-wise translations in the axial plane with spline registration. The resulting mean image of the motion corrected images was used for segmentation of the cord using *propseg* (manual correction was done when necessary). The mean moco image was warped into the PAM50 space using the concatenated warping field obtained at the anat step (i.e., from T1w to T2w to PAM50 space) to initialize the registration (with SCT, ‘*sct_register_multimodal’*).

As for the brain volumes, they undergo a standard preprocessing (Power et al. 2014, 2012) using FSL ‘*bet’* function for estimating the masks, FSL ‘*mcflirt’* to apply motion correction, and coregistering the functional images to T1w and the MNI template using SPM12.

For each participant, the nuisance regressors were modeled to account for physiological noise (Tapas PhysiO toolbox, an SP extension (Kasper et al., 2017)). Peripheral physiological recordings (heart rate and respiration) using the RETROspective Image CORrection (RETROICOR) procedure (Glover, 1999)). Specifically, four respiratory, three cardiac harmonics were modelled, and one multiplicative term for the interaction between the two (18 regressors in total, (Kinany et al., 2024; Landelle et al., 2024)). CompCor approach (Behzadi et al., 2007) was used to identify non-neural fluctuations in the unsmoothed brain or CSF signal by extracting the first 12 principal components for the brain and the first 5 for the spinal cord. These nuisance regressors were complemented with the two spinal cord motion parameters (x and y), the six brain motion parameters and motion outliers.

Similar to dataset 1 and 2 the removal of the noise confounds was based on a projection onto the orthogonal of the fMRI time-series space and was applied orthogonally to the band-pass temporal filter (0.01-0.13 Hz) using the Nilearn toolbox (‘*clean_img’* function (Lindquist et al. 2019)).

#### 2.3.4. Parcelled time series

##### Spinal cord

We defined a parcellation scheme consisting of 14 spinal cord regions in the cross-section characterized by 6 bilateral regions of the gray matter (intermediate zone (*iz*), ventral (*vh*) and dorsal horns (*dh*)) and 8 bilateral white matter regions (spinal lemniscus (SL), cortico-spinal tract (*cst*), fasciculus cuneatus (*fc*) and gracilis (*fg*)). These anatomical regions were delineated according to the atlas of the Spinal Cord Toolbox (De Leener et al. 2017). Since coherent activity has been reported in the white matter of the animal spinal cord (Paquette et al., 2021; Sengupta et al., 2021), and several studies have examined the reliability of functional signals in cerebral white matter (Gore et al., 2019; Wu et al., 2017), , we included spinal cord white matter ROIs to capture their potential role in brain–spinal interactions (particularly in *Dataset 3*), to ensure dimensional comparability with the brain parcellation, and to maintain consistent parcellations across datasets. To further mitigate partial volume effects, the SCT-derived WM ROIs were adjusted by removing bordering voxels, thereby reducing overlap with adjacent tissues. The number of spinal levels included is determined by the acquisition coverage (Figure 1) and the overlap among participants. Levels at the extremities were cropped in some participants; thus, to maintain consistency, only levels fully overlapping across all participants are considered. Specifically, *Dataset 1* included 3 spinal levels (C4–C6), resulting in a parcellation of 42 regions of interest (ROIs); *Dataset 2* comprised 5 fully overlapping spinal levels (C4–C8), yielding a dimensionality of 70 parcelled time series; and *Dataset 3* spanned spinal levels C2–C8, covering a total of 7 spinal levels and resulting in 98 ROIs. The voxel time series from the registered, denoised functional images were averaged using a robust mean, considering only time points between the 5th and 95th percentiles. In the paper we will refer to the number of spinal cord ROIs with 𝑁_!”_.

**Figure 1.**
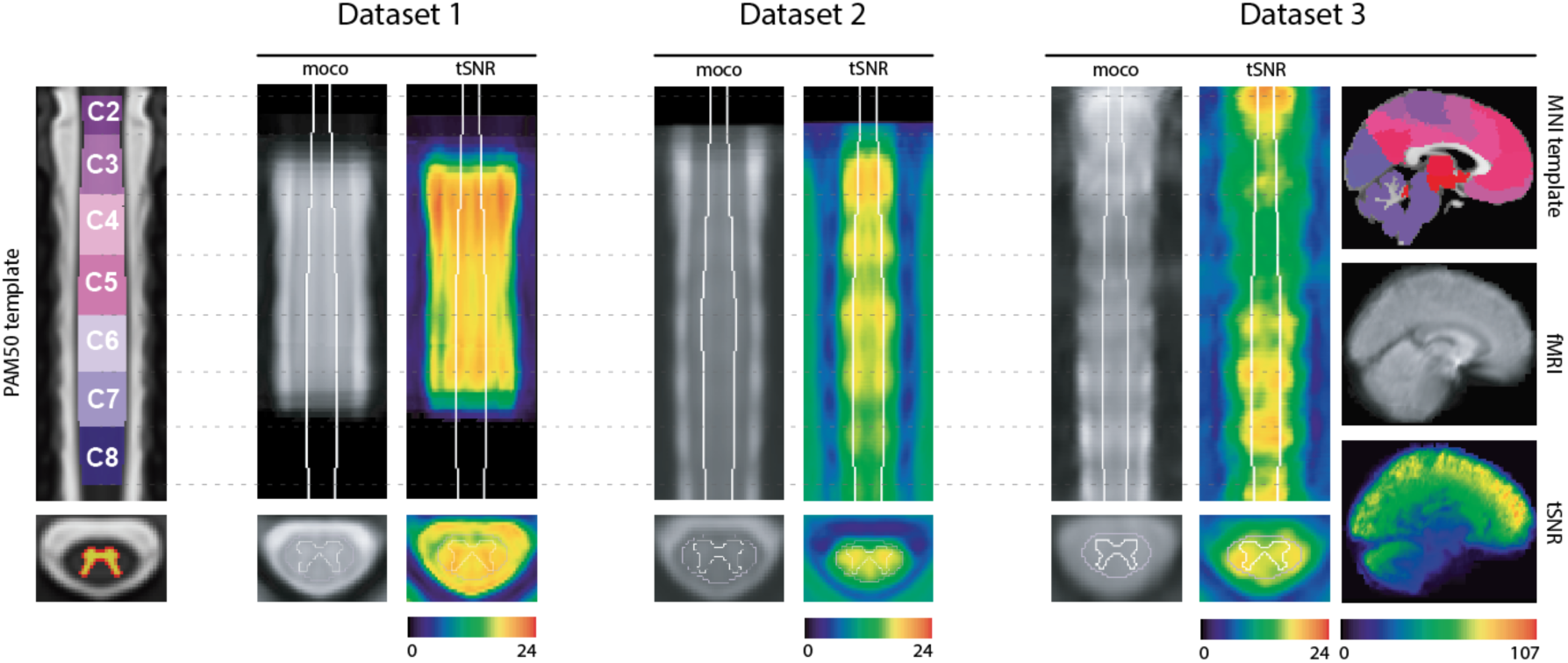
Datasets overview. Left panel: coronal view of the PAM50 template (y=70) with spinal level annotations (C2-C8) and the gray matter probability mask on the axial view. The three datasets are shown sequentially with the mean motion-corrected fMRI and the tSNR maps. *Dataset 1*: voxel size = 1 × 1 × 5 mm^3^, VAT = 2080 ms, protocol: 3D multi-shot gradient-echo sequence. *Dataset 2*: voxel size = 1.6 × 1.6 × 4 mm^3^, VAT = 2312 ms, protocol: 2D EPI. *Dataset 3*: voxel size = 1.6 × 1.6 × 4 mm^3^, VAT = 1550 ms, protocol: 2D EPI. *Dataset 3* additionally includes whole-brain coverage, with the brain data shown alongside the MNI template of 2mm resolution in the sagittal plane (x=45).

##### Brain

For *Dataset 3,* we also extracted parcelled brain time series following the same procedure: computing the robust mean across voxels within each of the 100 ROIs defined by the Schaefer’s atlas (Schaefer et al. 2018) and complemented by 19 subcortical areas (Seitzman et al. 2020), resulting with a parcellation of 𝑁_#$_ = 119.

### 2.4. Fingerprinting

#### 2.4.1. Methodology

We follow the steps proposed in earlier work at the brain level (Amico & Goñi, 2018). Let us assume 𝑁_!_ participants for which 𝐹*_i_^r^* denotes the FC matrix of run 𝑟 for participant 𝑖. Each matrix 𝐹*_i_ ^r^* has a dimensionality of 𝑁 × 𝑁, where 𝑁 indicates the number of ROIs (which can be 𝑁*_sc_* or 𝑁*_br_*, depending on whether spinal cord or brain regions are considered). The concept of fingerprinting centers on the ability to identify individuals based on these FC patterns; i.e., by comparing an individual’s FC matrix from one run against those from all participants, including another run from the same subject (Amico & Goñi, 2018; Finn et al., 2015). In this framework, two runs are available per subject (test-retest). To enable comparison, the upper triangular part of each 𝐹*_r_ ^r^* is unfolded to a vector 𝑐*_i_*^(*r*)^ of dimension 𝑀 = 𝑁(𝑁 − *1*)/*2*, to which Pearson correlation is applied. This results in the so-called identifiability matrix 𝐼 , of size 𝑁*_s_* × 𝑁*_s_* , which is asymmetric. The asymmetry arises because the correlation between run 1 of subject *i* and run 2 of subject *j* is not necessarily identical to the correlation between run 2 of subject *i* and run 1 of subject *j*. In other words, we are correlating across two ordered sets (run-1 → run-2), The diagonal elements 𝐼[𝑖, 𝑖] reflect self-similarity between the two runs of the same participant, while the off-diagonal elements 𝐼[𝑖, 𝑗] (with 𝑖 ≠ 𝑗) capture between-participant similarities. Let 𝐼*_self_* be the average of the diagonal elements, and 𝐼*_others_* the average of the cross-individuals similarities. We can now define a fingerprinting performance metric (*‘Idiff’*) as the difference between these values (𝐼*_self_*_+_ -𝐼*_others_*). Moreover, following also (Amico & Goñi, 2018), we explored the PCA of the FC profiles to maximize 𝐼𝑑𝑖𝑓𝑓 by applying dimensionality reduction (see supplementary material for details).

To further quantify the strength of the fingerprinting effect, we calculated Cohen’s D (Cohen 1992) as a standardized measure of the difference between two means (𝐼*_self_* and 𝐼*_others_* in particular), reflecting how large this difference is relative to the variability within the data. Let 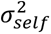 be the variance of the self-similarities and 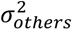 the variance of the off-diagonal elements, then the effect size is defined as follow:

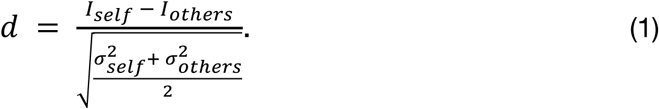

A higher Cohen’s D value indicates a greater separation (in standard deviation units) between individuals, thus stronger identifiability.

Following Finn et al., 2015, we also calculated the success rate as the number of participants that could be identified. A participant was considered correctly identified if the highest correlation in their row of the identifiability matrix appeared on the diagonal. This indicates that their FC profile from the first run was more similar to their own profile than to that of any other participant from the second run. We also propose a more lenient measure by computing the participants’ identification accuracy based on whether the correct participant was within the top 2, 3, 4, or 5 highest correlation values.

To assess which FC patterns contributed most reliably to individual identifiability, we used the one-way random model for the intraclass correlation coefficient that is known as ICC(1,1) (McGraw and Wong, 1996). Specifically, this metric measures the absolute agreement of repeated measurements for each connection across individuals:

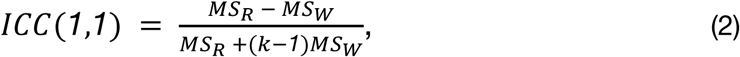

where 𝑘 = *2* (number of runs), 𝑀𝑆_0_ is the mean square between individuals, and 𝑀𝑆_1_ is the within-participant (error) variance. This computation was repeated independently for each edge, yielding a vector of ICC values, which were then mapped back to the 𝑁 × 𝑁space to obtain the full ICC matrix. This method provides a fine-grained identifiability map of which FC edges are most stable across runs, reflecting their test-retest reliability. To have an overall overview of these reliable edges in the context of the spinal cord, we averaged the ICC matrices across datasets and all levels. For the cervical spinal cord, we recall that the number of ROIs is 14 in the cross-section (i.e., bilaterally for gray matter: *dh, vh, iz;* and for white matter: *cst, fc, fg, sl*). Moreover, we examined the ICC matrices after thresholding at the 95th percentile to retain only the most reliable edges. To shift the focus from the connections to region-based reliability, we computed the nodal strength of this filtered matrix by summing the values across one axis, providing a measure of reliability for each ROIs rather than for specific connections.

#### 2.4.2. Investigating brain and spinal cord time courses

To further explore Dataset 3, we applied the fingerprinting pipeline to the full FOV including brain and cervical spinal cord, and looked at three FC profiles: brain-only, spine-only, and brain-spine interaction. In addition, we examined the relationship between brain and spinal time series using a simple linear regression framework (equation 3), with the goal of removing shared variance. Since the imaging was acquired simultaneously, the time axis is shared between the dependent variable (𝑌) and the independent variable (𝑋), both of dimension 𝑁 × 𝑇, where 𝑁 is the number of ROIs (spinal or brain) and 𝑇 the number of time points. The regression model is defined as:

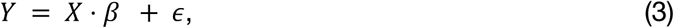

where Ŷ = 𝑋 ⋅ 𝛽 are the predicted values and 𝜖 = 𝑌 − Ŷ represents the residuals.

This framework was then applied in two directions:

1. Spine residuals (brain regressed out):

○ Dependent variable: 𝑌 = 𝑌*_sc_* (spinal ROIs).

○ Predictors: 𝑋 = 𝑋*_br_* (all brain ROIs).

○ Predicted spinal activity: 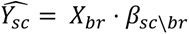.

○ Residuals: 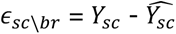 capturing spinal variance not explained by brain signals.

○ Dimensionality: 𝑁*_sc_* × 𝑇.

2. Brain residuals (spine regressed out):

○ Dependent variable: 𝑌 = 𝑌*_br_* (brain ROIs).

○ Predictors: 𝑋 = 𝑋*_sc_* (all spinal ROIs).

○ Predicted brain activity: 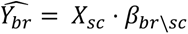.

○ Residuals: 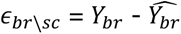 capturing spinal variance not explained by brain signals.

○ Dimensionality: 𝑁*_br_* × 𝑇.

In summary, the first model uses all brain regions to predict spinal activity, whereas the second uses all spinal regions to predict brain activity.

We then used the residuals to compute new FC profiles that reflect variance not accounted for by the other region.

## Results

### 3.1 Data quality and overview

In Figure 1, we present an overview of the three datasets analyzed in this study together with the template spaces that were employed: PAM50 for spinal cord data (left) and MNI for brain data (right, shown alongside *Dataset 3*, which covers the brain). The ROIs in the brain were defined using the Schaefer-100, complemented by 19 subcortical areas, and are visualized on the MNI template. For the spinal cord, only spinal levels C2-C8 are shown on the PAM50 template, reflecting the coverage range of the spinal cord acquisitions across datasets. Mean fMRI images and tSNR maps of the first run are also provided for each dataset. In the spinal cord, tSNR values reach a maximum mean value of 22.6 in *Dataset 3*, followed by 21.3 for *Datasets 1*, while in *Dataset 2,* the maximum value is 19 (within the cord mask). By comparison, brain tSNR values are substantially higher, reaching up to 107.

### 3.2. Spine-print: fingerprint of the spinal cord

In Figure 2, we display the FC (A) and identifiability matrices (B) derived from the three datasets with a focus on the spinal cord only. The identifiability matrices are reordered by diagonal correlation values in descending order. The zoomed-in panels in the first column highlight the anatomical organization of the ROIs in the spinal cord, with gray and white matter regions alternating from left to right—a sorting scheme that remains consistent throughout the paper whenever spinal levels are referenced. Across all datasets, the strongest correlations were observed ipsilaterally, both in gray and white matter, most likely reflecting spatial proximity and partial volume effects (e.g., between the fasciculus gracilis and the adjacent fasciculus cuneatus on the same side). Beyond these local effects, consistent bilateral (left–right) correlations were also detected. The most prominent of these involved the **intermediate zones (*iz*)**, followed by the **ventral horns *(vh)***, the **fasciculus cuneatus (*fc*)**, the f**asciculus gracilis (*fg*)**, and, lastly, the **dorsal horns *(dh)*.**

**Figure 2.**
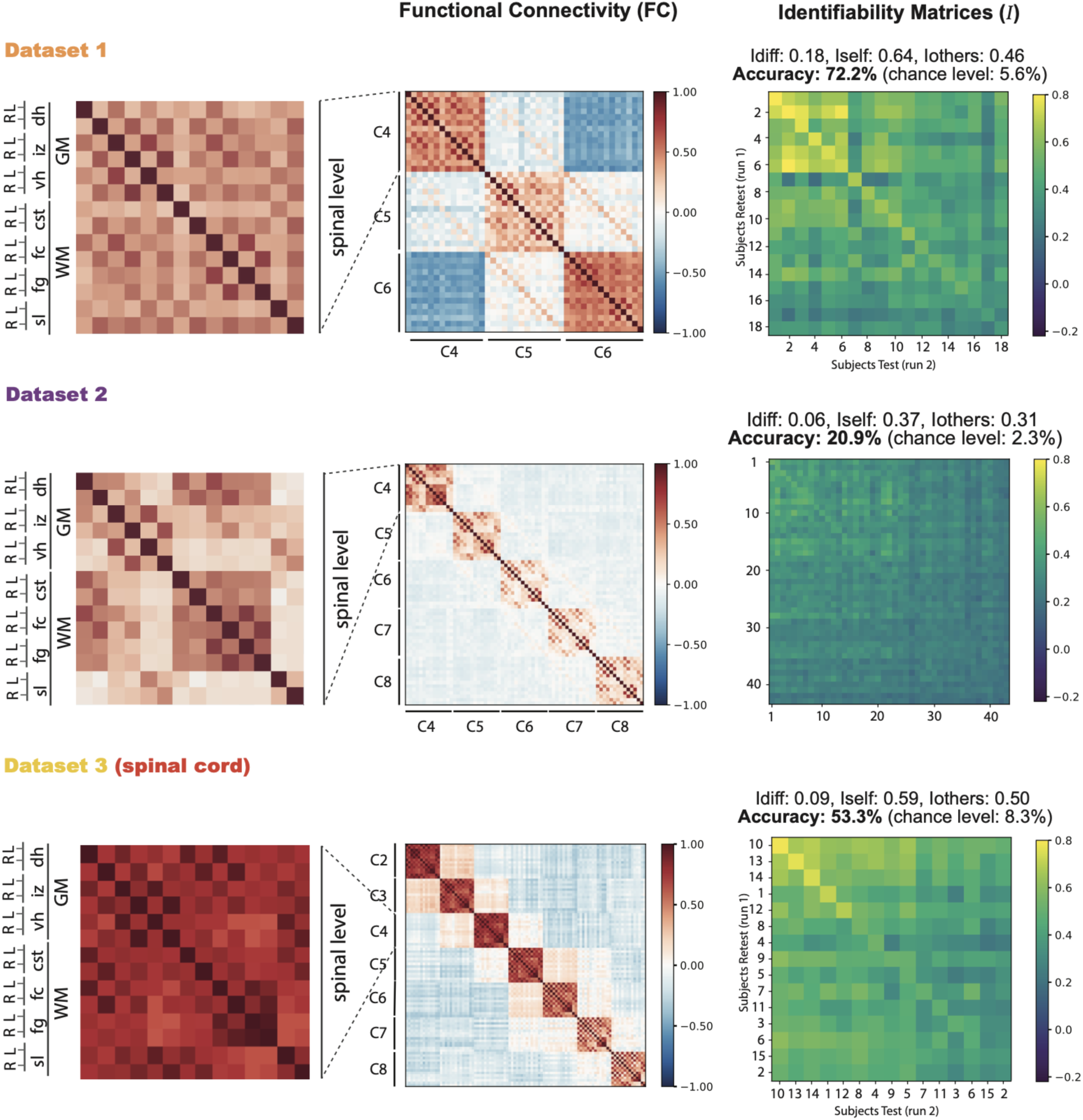
Spine-print. Overview of functional connectivity (FC) matrices and identifiability results for the spinal cord across the three datasets. FC matrices of the first run from each dataset. The most-left matrix is a zoomed-in view of the spinal level C4 for all datasets to show the labelling and sorting of the spinal cord ROIs used for all the spinal levels in all matrices. B. The third column shows the corresponding identifiability matrices for each dataset, along with the reached scores and accuracies. Matrices are reordered based on diagonal correlation values. For comparison with Figure 3, subject IDs are reported only for Dataset 3, where the ordering is relevant.

In Dataset 1, the strongest correlations were observed ipsilaterally between the *fg* and *fc*, reaching 0.51 on the right and 0.50 on the left. These were followed by gray matter connections between the *iz* and the *vh* (0.50 right, 0.49 left). Next, the *dh* connected with the *iz* (0.38 right, 0.34 left), while bilateral correlations within the ventral horns reached 0.29 and within the *fc* 0.26. The *fg* showed weaker bilateral correlations (0.24), and the lowest among the main gray matter regions was observed within the dorsal horns (DL-DR) with 0.17.

*Dataset 2* shows a similar profile. The strongest correlations were again ipsilateral between the *fg* and *fc*, reaching 0.50 on the right and 0.50 on the left. Among gray matter regions, the *iz* connected strongly with the *vh* (0.42 left, 0.41 right). The *dh* also showed notable ipsilateral correlations with the *fc* (0.35 left, 0.31 right). Further gray matter interactions included *vh*–*iz* correlations (0.28 right, 0.27 left). Bilateral correlations were overall weaker, with the *fg* reaching 0.28 and the *vh* 0.13, while the *dh* showed the lowest bilateral connectivity (0.12).

In Dataset 3, overall correlations were stronger than in the other datasets, preserving similar patterns. The highest value was observed for bilateral connectivity of the *fg* (0.82). This was followed by gray matter connections between the *iz* and *vh* (0.80 left, 0.80 right) and strong ipsilateral correlations between the *dh* and *fc* (0.80 left, 0.80 right). Bilateral *fg*–*fc* connectivity also remained high (0.75 left, 0.74 right). The **corticospinal tract (*cst*)** showed robust correlations with the *dh* (0.69 right, 0.68 left). Within the gray matter, *dh-iz* correlations reached 0.61, while *dh-vh* connectivity followed at 0.49. Finally, bilateral correlations in the *vh* (VL-VR: 0.33) and *dh* (DL-DR: 0.27) were weaker but consistent across levels.

To assess the intersubject reliability of these averaged correlations, we computed the identifiability matrix (𝐼), which quantifies the extent to which each participant’s FC profile is distinct within the dataset across two runs (Figure 2). We recall that 𝐼 is an 𝑁_!_ × 𝑁_!_ matrix, where 𝑁_!_ is the number of participants in a dataset. These matrices exhibit a clear diagonal pattern, indicating that each participant’s FC profile is more similar to their own than to anyone else’s. The maximum correlation values observed on 𝐼 are 0.85 for *Dataset 1*, 0.6 for *Dataset 2*, and 0.79 for *Dataset 3*. *Dataset 1* reaches an 𝐼𝑑𝑖𝑓𝑓 = 0.18 with a subject identification accuracy of 72.2% (13 participants were correctly identified over 18, chance level = 5.6%). *Dataset 2* reaches an 𝐼𝑑𝑖𝑓𝑓 = 0.06, accuracy = 20.9% (9 participants correctly identified over 43, chance level = 2.3%). *Dataset 3* 𝐼𝑑𝑖𝑓𝑓 = 0.09, accuracy = 53.3% (8 / 15 correctly identified, chance level = 8.3%). Cohen’s D effect sizes of *Iself* and *Iothers* provide a measure of the magnitude of their difference. That is, *Datasets 1 and 3* exhibit very large effect sizes with 1.84 and 0.87, respectively. Despite the relatively low 𝐼𝑑𝑖𝑓𝑓 values and accuracy in *Dataset 2,* it still shows a medium effect size (Cohen’s D = 0.71). Typically, effect sizes are considered small between 0.2–0.4, medium between 0.5–0.8, and large above 0.8.

Having characterized the spine-print, we next focus on *Dataset 3* as a whole, examining not only spinal signatures but also brain connectivity and its interaction with the spine (Figure 3A-B).

**Figure 3.**
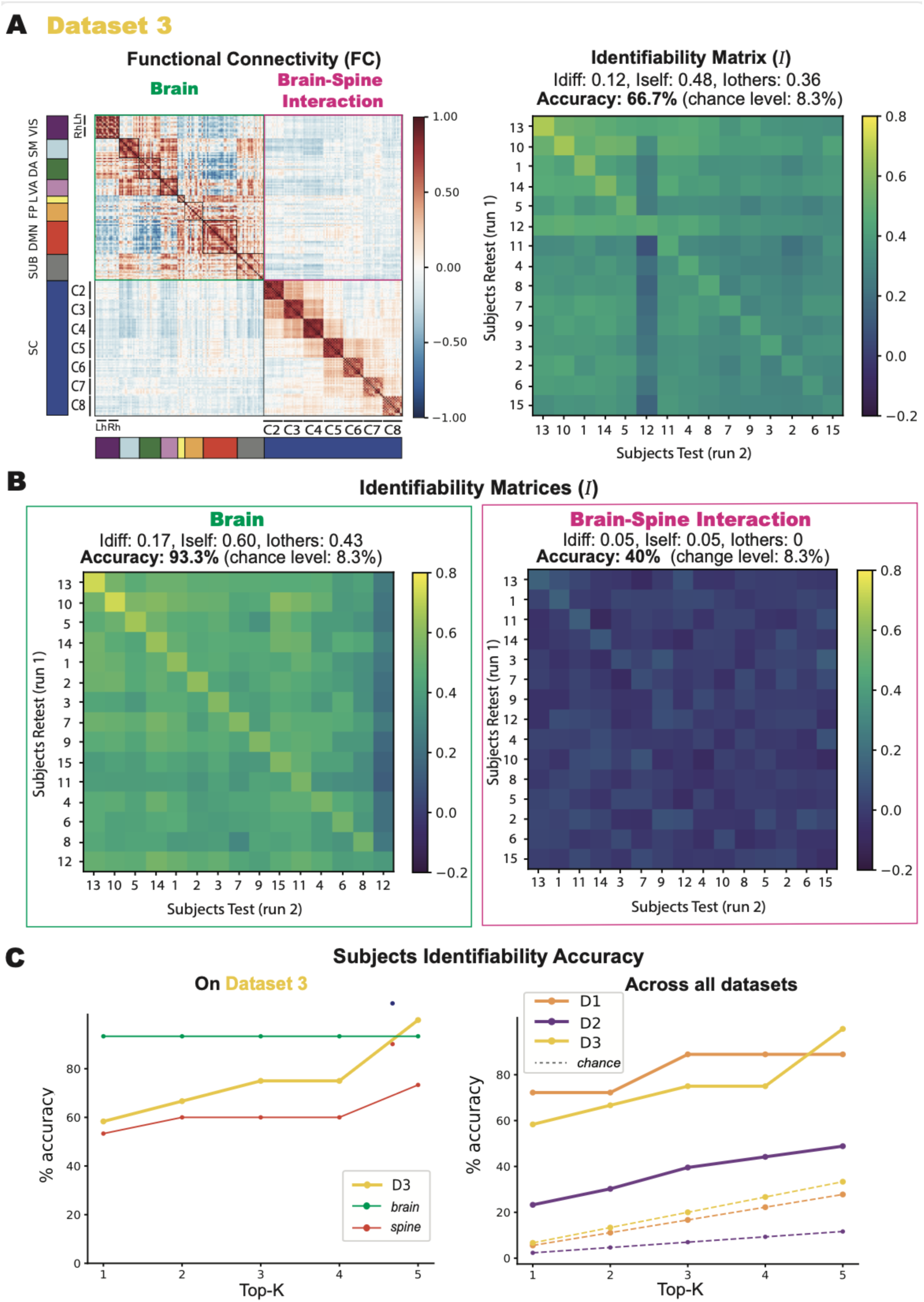
Dataset 3 fingerprinting. The identifiability matrices reflect the correlations across the participants’ runs, with participants’ IDs sorted by highest correlation in each case, leading to differing axis labels across matrices. Identification accuracies (chance level = 8.3%) are 93.3% for brain-only, 53.3% for spine-only, and 40% for the interaction. (C) On the left, a plot illustrates participants identifiability accuracy as a function of the top-K (from 1 to 5) highest correlation values with a focus on *Dataset 3*, with separate curves for brain-only and spine-only data. On the right, the same plot across the three datasets.

**Figure 4.**
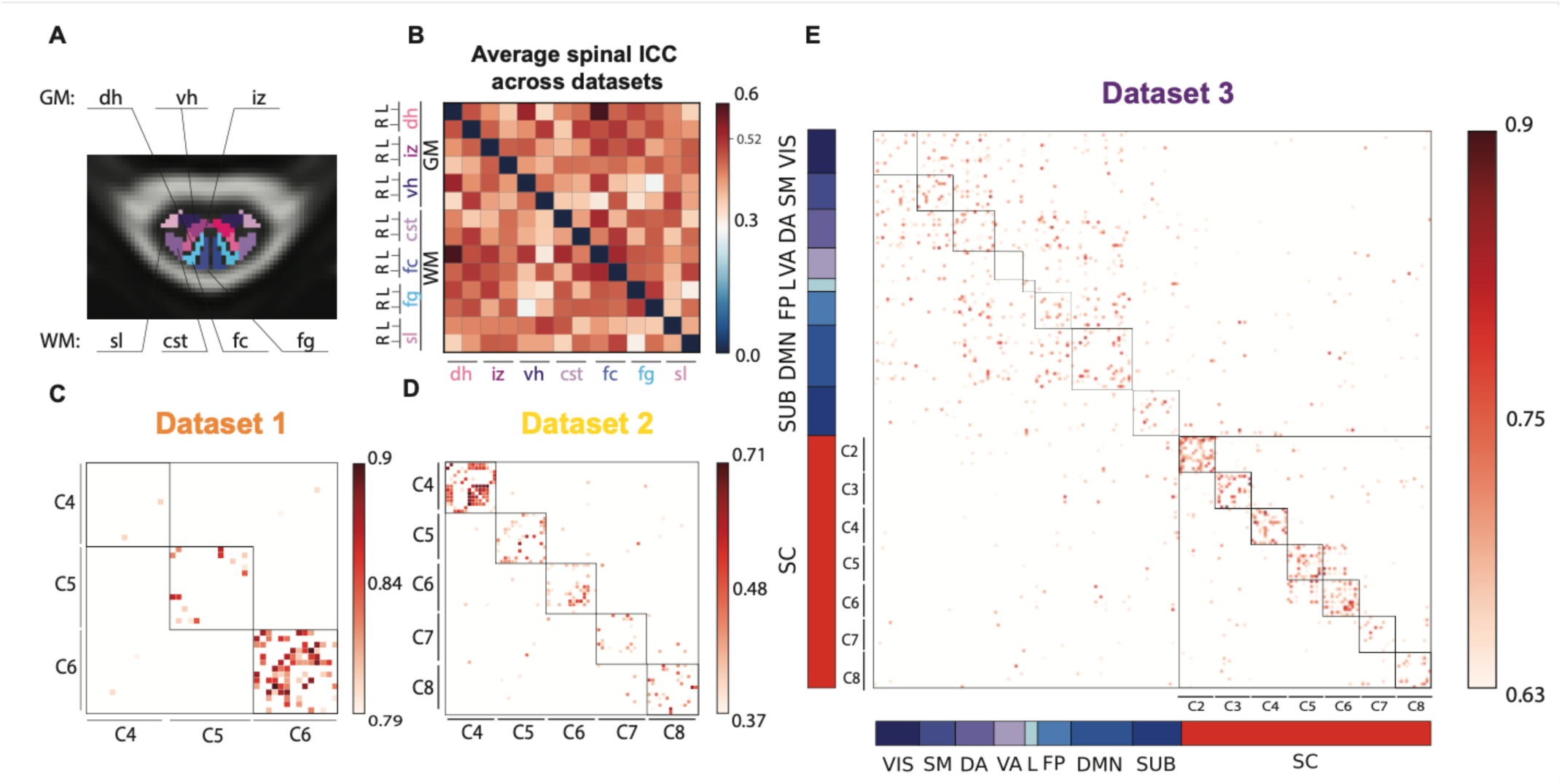
ICC results. (A) Spinal cord ROIs displayed in a cross section of the spinal cord (gray matter (GM) regions: *dh* = dorsal horns, *iz* = intermediate zone, *vh* = ventral horns, white matter (WM) regions: *sl* = spinal lemniscus, *cst* = cortico-spinal tract, *fc* = fasciculus cuneatus, *fg* = fasciculus gracilis). (B) Average ICC matrix across all spinal levels and datasets, indicating the 95th percentile in the colorbar (0.526). (C-D-E), namely, Datasets 1,2, and 3. (E) For *Dataset 3*, brain ROIs are sorted according to the Yeo functional networks (Yeo et al. 2011) (VIS = visual, SM = somatomotor, DA = dorsal attention, VA = ventral attention, L = limbic, FP = fronto-parietal, DMN = default mode network, SUB = subcortical). The color bar values are adjusted according to the distribution of the values, reporting the 95th percentiles of each dataset’s ICC as minimum to show a filtered ICC matrix.

### 3.3. Fingerprint of the central nervous system

We conducted a detailed analysis of Dataset 3, examining functional connectivity within the brain, within the spinal cord, and across brain–spinal interactions. Figure 3A shows the full FC matrix (left) and the corresponding identifiability matrix (right). Brain regions were ordered according to Yeo’s functional networks (Yeo et al., 2011), concatenating left and right hemispheres within each network. For each component (brain, spine, brain–spine), identifiability matrices are shown alongside their corresponding accuracy and identifiability scores. The full dataset achieved an accuracy of 66.7% (8/15 individuals), an improvement of two correctly identified participants compared with the spine-only matrix.

As shown in Figure 3B, the brain-only matrix (highlighted in green) yielded the highest accuracy at 93.3%, with 14 of 15 individuals correctly identified. By contrast, the brain–spinal interaction (in pink) produced the lowest accuracy of 40% (6/15 individuals). Effect sizes were very large for the full dataset (Cohen’s d = 1.43) and brain-only (d = 1.98), clearly exceeding those for the spine-only (d = 0.96) and brain–spine (d = 0.83) matrices. Participant IDs were sorted independently within each component by decreasing diagonal correlation values, and therefore do not align across matrices. Still, participants 1, 10, 11, and 13 consistently appeared among the top of the matrices in all conditions: spine-only (Figure 2), the full dataset (Figure 3A), and the brain-and brain–spine–only analyses (Figure 3B, respectively in green and pink). All three matrices showed a visible diagonal pattern of identifiability, most pronounced in the brain matrix. Additional details are provided in the Supplementary Material.

To further explore accuracy, Figure 3C plots performance as a function of the top-K best correlations. For *Dataset 3 (*left column), brain accuracy remained constant across K, whereas spine accuracy increased when more top correlations were considered. The right column shows that this upward trend held consistently across all three datasets.

Finally, we examined the residual correlations from linear regression models to assess their impact on subject identifiability. In the first model, where spinal cord time series served as the dependent variable (𝑌) and brain time series as the predictor (𝑋), the residuals— representing spinal cord activity not linearly explained by brain activity—yielded an 𝐼𝑑𝑖𝑓𝑓 score of 0.07, with 7 out of 15 participants correctly identified (accuracy = 46.7%). Conversely, in the inverse model, with brain time series as 𝑌 and spinal cord time series as 𝑋, the residuals—indicating brain activity unexplained by spinal cord signals—produced a higher 𝐼𝑑𝑖𝑓𝑓 of 0.12 and a similar identification, correctly identifying 8 out of 15 individuals (accuracy = 53.3%).

### 3.4. Reliability of individual connections in functional fingerprinting

The intraclass correlation coefficient (ICC) was used to evaluate the reliability of FC patterns across the two runs. To summarize spinal cord ROI reliability, we computed the mean ICC values across cervical levels for the three datasets (Figure 3B). According to standard guidelines (Hallgren 2012; Cicchetti and Sparrow 1981), ICC values are interpreted as: poor (<0.4), fair (0.4–0.59), good (0.6–0.74), and excellent (≥0.75).

Across all datasets, the most reliable connection in the spinal cord—above the 95th percentile threshold (ICC = 0.52)—was observed between the left dorsal GM horn and the left fasciculus cuneatus (ICC = 0.596), followed by ventral GM horns and dorsal GM horns (ICC = 0.54), both falling within the “fair” reliability range.

Looking at each dataset individually: (i) *Dataset 1* showed excellent ICC values, particularly at the C6 level. When averaging ICC values across the three spinal levels to obtain nodal strength per ROI, the left *fc-dh* connectivity had the highest average ICC (0.81), followed by the bilateral *fc* with 0.78, (Figure 3C). *Dataset 2* exhibited overall lower ICC values, with the 95th percentile at 0.41 and a maximum ICC of 0.45 (interpreted as “fair”). Here too, nodal strength was highest for the left *fc-dh* connectivity (0.45), followed by the *fg*-*dh* on the left side, and lowest for the bilateral *cst-fc* (0.41) (Figure 3D). *Dataset 3* had higher reliability overall, with ICC values above 0.57 (95th percentile), and a maximum ICC of 0.86 (Figure 3E). In the brain, the most reliable connection (0.86) was the left salient ventral attention (VA) with the left DMN prefrontal cortex 5 (DMN) and the lowest ICC reached 0.63 with the left visual network 9 (VIS) and the right somatomotor 8. In the spinal cord, the most reliable connection was right *fc* with right spinal lemniscus (*sl*) with a good ICC of 0.6 and the lowest (0.57) was between *fc* and *vh* right.

Considering the nodal strength as mean values across the ICC matrix along one axis, in the brain, the values were relatively consistent across regions, averaging around 0.68, with the visual network and subcortical regions showing the lowest value (0.64), and default mode network (DMN) and limbic network (L) with 0.7. Interestingly in all three datasets, the spinal cord exhibits the strongest nodal strength for fasciculus cuneatus (*fc*), followed by fasciculus gracilis (*fg*) and dorsal horns (*dh*), while the lowest values differ in the three datasets: *Dataset 1* smallest nodal strength was found in the ventral horns, in *Dataset 2* it was the intermediate zone, and the *cst* for *Dataset 3*.

## 3. Discussion

Our findings demonstrate that a reliable and individually distinctive FC pattern from the (cervical) spinal cord – termed “spine-print” – can be identified across individuals in three different datasets. Despite differences in acquisition protocols and slight variations in preprocessing, participants’ identification remains well above chance, highlighting the feasibility and robustness of the spine-print. The different performances across the three datasets may partly stem from differences in acquisition type. In particular, the dataset showing the best performance (*Dataset 1*) was acquired with 3D EPI, which is known to yield higher tSNR (Lutti et al., 2013), whereas the other two relied on 2D EPI. Beyond this, the lower absolute scores observed for *Dataset 2* compared to the others may also reflect the longer time interval between its two runs, which could reduce within-subject consistency. Although manual versus automatic shimming might influence results, tSNR did not differ significantly between runs (Kaptan et al., 2022, 2023), suggesting that the effect cannot be explained solely by signal quality. Importantly, despite these differences, identifiability in *Dataset 2* remained nearly an order of magnitude above chance, underscoring the robustness of the approach even under less favorable conditions. Overall, much like the brain, the spinal cord exhibits participant-specific FC that is reliably detectable.

A closer inspection of the spine-prints and the regions with the highest reliability reveals that the most consistent areas—those with the highest nodal strength—were the sensory regions, rather than the ventral regions where motor neurons reside. This could suggest that the sensory functions processed by the cord might be more subject-specific, while motor functions at rest are more uniform across individuals. In *Dataset 3*, we also assessed the identifiability scores of the sensory-motor (SM) network alone in relation to the spinal cord, allowing for a fair comparison between the two. Interestingly, when the SM network and spinal cord regions were combined, the identifiability scores increased notably (see Supplementary Material, Fig. S4).

In terms of correlations, the ipsilateral connectivity of the fasciculus gracilis (*fg*) with fasciculus cuneatus (*fc*) consistently emerged as the strongest across all three datasets. The dorsal cord is particularly sensitive to physiological artefacts such as CSF pulsatility and cardiac-driven motion (Brooks et al., 2008). Because *fg* and *fc* are adjacent dorsal column tracts, they may be affected in phase by the same physiological noise sources. Both convey somatosensory information (*fg* = lower body, *fc* = upper body) (Yoganandan et al., 2017), and during rest, ongoing sensory afferents could still drive synchronous baseline fluctuations across these pathways. However, such functional explanations are more difficult to establish and are likely secondary to spatial proximity and shared vascular supply. Indeed, both tracts are perfused by the posterior spinal arteries, and the most plausible explanation for their strong correlations remains partial volume effects arising from their anatomical adjacency.

In *Dataset 3* the bilateral connection of *fg* was the strongest. Yet it is important to note that this dataset was acquired at a lower resolution, making it more prone to inflated correlation values among anatomically neighbouring ROIs. More generally, many of the strong bilateral correlations observed across datasets could be largely influenced by partial volume effects, given the small size and close proximity of these regions.

When assessing edge reliability with ICC, we found that the connection between the fasciculus cuneatus (*fc*) and the dorsal horns (*dh*) showed the highest score. We note, however, that the close anatomical proximity of these regions means partial volume effects could contribute to this result. At the nodal level, reliability was highest for sensory regions (*fc, fg, dh*), followed by motor regions (*cst, vh, iz, sl*). This pattern may indicate that, in the absence of an explicit task, the ventral horn exhibits greater variability in its spontaneous signal, whereas the signal in the dorsal horn remains more stable to maintain consistent sensory processing for body awareness at rest.

Additionally, the findings in the dorsal areas could also be affected by the presence of the dorsal vein, potentially confounding the interpretation of BOLD signals in these regions. While the interpretation of BOLD signals in the white matter remains debated, there is growing evidence that white matter has sufficient vascularization to support hemodynamic changes. Several studies have investigated the reliability of functional information captured in the cerebral white matter (Wu et al., 2017; Peer et al., 2017; Huang et al., 2020; Ding et al., 2018; Gawryluk et al., 2014; Gore et al., 2019). White matter is largely composed of glial cells, which regulate homeostasis and support myelinated axons, and their metabolic processes can influence local blood flow and presumably influence the oxygenation demand of the WM. These non-neuronal fluctuations may explain the distinct slower dynamics of BOLD signals in white matter compared to gray matter. Recognizing these factors is essential for accurate interpretation of spinal cord fMRI results (Paquette et al., 2021; Sengupta et al., 2024). In line with this, dynamic functional connectivity approaches have revealed functional components in the spinal cord white matter during rest, as shown by Kinany et al., 2020.

Importantly, an individual-specific signature could be primarily shaped by anatomical features. Prior studies have shown that spinal cord functional patterns tend to follow its anatomical segmentation—for example, localizing in line with the defined dorsal/ventral horns as well as white-matter tracts (Kinany et al., 2020; Ricchi, Gomez, et al., 2024; Sengupta et al., 2021) supporting the notion that inter-individual anatomical variability may give rise to detectable functional differences.

Regardless of the potential functional or anatomical nature of the spine-print ,, its distinctiveness is notably weaker compared to that of the brain. This difference raises important questions about the factors influencing the strength of individual signatures in different parts of the central nervous system, which we discuss next.

One such factor could be the relative granularity of the parcellation used to define ROIs. Notably, the number of ROIs in Dataset 3 was kept relatively comparable between the brain and the spinal cord: the brain was divided into 119 ROIs, while the spinal cord included 98, derived from 14 cross-sectional regions across 7 spinal levels. Yet, despite this similarity in dimensionality, the physical sizes of these structures differ drastically (Parent & Carpenter, 1996; Sherman et al., 1990). Their volumes are 1200-1400 cm^3^ and about 11 cm^3^, respectively, considering 8-10 cm of length for C1-C7 and a cross-section area of 1-1.5 cm^2^. This results in a volume ratio of ∼118. The ROIs from the atlases used for each are on average ∼227 times smaller for the spine than for the brain. As a result, regional signal averages in the spinal cord are expected to be inherently noisier, especially given the lower tSNR typically observed in spinal fMRI. These factors together help explain the comparatively lower fingerprinting performance observed in the spinal cord relative to the brain. Thus, even though identifiability scores appear lower when compared directly to the brain, it is noteworthy that reliable spine-prints could still be detected despite spinal tSNR being roughly five times lower than in the brain (see Figure 1). At the same time, an important limitation of fingerprinting approaches is that they cannot fully disentangle neurobiological features from individually stable artefacts. Subject-specific factors such as susceptibility-related signal dropout, residual motion, or acquisition-related effects may remain consistent across sessions and thus contribute to identifiability without necessarily reflecting neural processes. Similar concerns have been raised in the brain fingerprinting literature (Amico & Goñi, 2018b; Bijsterbosch et al., 2018; Finn & Todd Constable, 2016; Horien et al., 2019; Xifra-Porxas et al., 2021), where fingerprint accuracy is acknowledged to reflect a mixture of neural and non-neural sources.Beyond size and signal quality, brain and spine also have different functional roles. For brain fingerprinting, the regions that are most specific for an individual are those serving higher-order cognitive functions, while sensory regions provide less unique information (Amico & Goñi, 2018; Finn et al., 2015; Griffa et al., 2022; Jo et al., 2021; Lu et al., 2024). The latter are evolutionarily more preserved, and, given that the spinal cord is an even more ancient structure, it might therefore be presumed to exhibit limited individual variability and, consequently, a diminished capacity for supporting a distinct “spine-print”. Yet, despite these expectations, our results reveal a measurable degree of individual specificity in spinal cord FC. This is supported by our regression analyses examining the linear relationship between brain and spinal cord temporal dynamics, thus investigating the potential impact of brain signals on spinal cord activity and *vice versa*. When cortical contributions were regressed out of the spinal cord time courses—leaving residuals for fingerprinting—the identifiability score decreased from 0.09 to 0.07, and participant identification remained at 7 out of 15. In contrast, regressing out spinal cord activity from the brain signals had a stronger effect: the identifiability score (𝐼𝑑𝑖𝑓𝑓) dropped from 0.17 to 0.12, matching the value obtained from the full brain+spine dataset. Participant identification in this case was 8 out of 15, still markedly lower than the original brain-only accuracy of 93.3% (14/15). These findings indicate that the existence of a “spine-print” cannot be solely attributed to brain time series; rather, spinal time courses exert a linear influence that enhances the identifiability of brain ROIs. Nonetheless, these conclusions rest on a linear regression framework, which assumes linear dependencies.

Together, these findings indicate that FC connectivity influences the correlation values of the identifiability matrix, without affecting the identifiability accuracy. While 𝐼𝑑𝑖𝑓𝑓 quantifies the *magnitude* of the separation between within- and between-subject similarity (i.e., how strongly self-similarity exceeds other-similarity), *accuracy* only captures the *outcome* of this comparison (i.e., whether self-similarity is higher than any other). Consequently, a low 𝐼𝑑𝑖𝑓𝑓 can still coincide with high accuracy, as even small margins are sufficient for correct identification as long as the self-similarity remains the top match.

Given the relatively small difference between the spine-only and spine-residual models, we avoid drawing conclusions that the brain primarily drives spinal cord activity. Furthermore, linear regression offers only a simplified perspective on dependency and cannot detect nonlinear or more complex bidirectional relationships, so these findings should be interpreted cautiously. Our results thus reveal some effects of brain and spine connectivity on identifiability, but future research employing more sophisticated modelling methods will be required to separate the relative contributions of signals from these two structures.

## 4. Conclusion and outlook

In this study, we demonstrate the feasibility of identifying an individual-specific functional signature in the spinal cord—a “spine-print”—across two runs within the same scanning session. This finding supports the notion that participant-level identifiability is not exclusive to the brain and can extend to spinal cord activity. Interestingly, brain regions involved in sensorimotor functions—those more directly related to spinal cord activity—tend to exhibit lower identifiability scores compared to higher-order cognitive regions such as the prefrontal and frontoparietal cortex (Amico & Goñi, 2018; Finn et al., 2015). This parallel may indicate that the reduced spine-print identifiability, relative to brain fingerprinting, reflects functional characteristics of sensorimotor systems more generally.

Future work should aim to explore the stability of this spine-print across sessions to determine whether it reflects a short-term phenomenon or a more persistent individual trait. In particular, systematically varying the time interval between consecutive runs or sessions would help assess whether identifiability decays over time, thereby clarifying whether the spine-print represents an instantaneous state-like signature or a more stable individual trait. However, this remains technically challenging. The spinal cord’s curvature is highly influenced by participant positioning and can vary significantly across sessions, posing challenges for accurate between-session image co-registration (Dabbagh et al., 2024). To address this, an informative next step would involve scanning the same individual in two consecutive sessions with only a brief pause in between, during which the participant is temporarily removed from the scanner but remains positioned on the table—compared to a condition where the participant is repositioned entirely. This approach would allow for a more controlled assessment of acquisition-related variability, disentangled from repositioning effects.

Additionally, our findings highlight the potential role of the brain in shaping spinal cord activity. Given the observed interactions, future research should further investigate the dependency and directionality of brain–spine functional connectivity, which may yield new insights into hierarchical control mechanisms within the central nervous system.

## Acknowledgments

Data acquisition for *Dataset 1* was primarily supported by National Institutes of Health (NIH) grants K99EB016689 and R21NS081437. Data processing was supported by NIH grants R00EB016689, R01EB027779 and R21EB031211. The content is solely the responsibility of the authors and does not necessarily represent the official views of the NIH.

Since January 2024, Dr. Barry has been employed by the National Institute of Biomedical Imaging and Bioengineering at the NIH. This article was co-authored by Robert Barry in his personal capacity. The opinions expressed in the article are his own and do not necessarily reflect the views of the NIH, the Department of Health and Human Services, or the United States government.

This work was supported by the Swiss National Science Foundation (SNSF), Project No. 205321_207493.

## Data and Code Availability

*Dataset 2* is publicly available, while *Dataset 1* and *3* are available upon reasonable request to Dr. Robert Barry and Prof. Julien Doyon, respectively.

Code is publicly available on the GitHub repository: https://github.com/MIPLabCH/SpinePrint.

## Authors contributions

I.R. formal analysis, investigation, conceptualization, methodology, software, validation, visualization, writing – original draft; A.S. methodology, supervision, writing – review & editing;

N.K. conceptualization, supervision, writing – original draft, review & editing; C.L. *Dataset 3* curation, writing – original draft, review & editing; A. K. and S. V. acquired *Dataset 3;* J. D. is the *Dataset 3* owner, writing – original draft, review & editing; R.B. owns *Dataset 1*, supervision, writing – original draft, review & editing; D.V.d.V. conceptualization, supervision, writing – original draft, review & editing.

## LIST OF ABBREVIATIONS

FC: functional connectivity
ROI: region of interest
ICC: inter cluster coefficient
Idiff: identifiability score
dh: dorsal horns
Iz: intermediate zone
vh: ventral horns
cst: cortico-spinal tract
fc: fasciculus cuneatus
fg: fasciculus gracilis
sl: spinal lemniscus
VIS: visual network
SM: somatomotor network
DA: dorsal-attention network
VA: ventral-attention network
L: limbic network
FP: fronto-parietal network
DMN: default-mode network
GM: gray matter
WM: white matter

## Notes

https://openneuro.org/datasets/ds004386/versions/1.1.2

https://www.ismrm.org/21/program-files/TeaserSlides/TeasersPresentations/0654-Teaser.html

### Competing Interest Statement

The authors have declared no competing interest.

### Summary of Updates

1) Figures order and organization. Figure 2 now contains the spine-print results, also for Dataset 3, while the brain-spine analysis has been moved to a separate figure; 2) New spine parcellation; 3) Conclusion on the regression analysis; 4) Additional figure in supplementary

