## supplementary material for "Spine-Prints: Transposing Brain Fingerprints to the Spinal Cord"

### Principal Component Analysis (PCA)

Building on the method proposed by Amico et al. (2018), we optimized the identifiability score (*Idiff*) using PCA. Specifically, we report the variance explained by each Principal Component (PC) and the cumulative distribution in the scree plots shown in Figure S1.

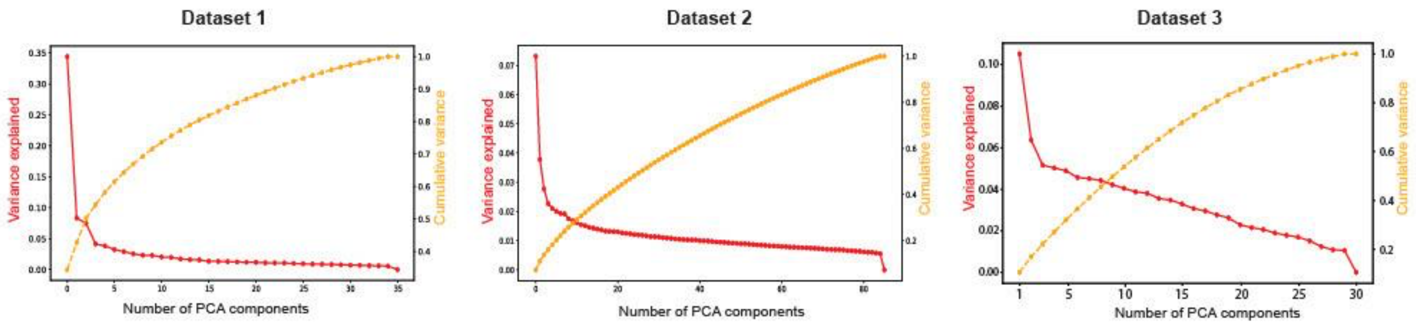


**Figure S1.** Scree plot of the variance explained (in red) and the cumulative variance (in yellow) as a function of the number of components across the three datasets.

We further examine how the $Idiff$ score (in red) and participant identification accuracy (in blue) vary as a function of the number of PCs, as illustrated in Figure S2.
The numbers of PCs yielding the highest $Idiff$ were 12 for *Dataset 1*, and for both *Dataset 2* and *3* the first 10 PCs were the ones that optimize both $Idiff$ and accuracy (see curves peaks in Figure S2). As discussed in Amico et al. 2018, there are as many components as the number of functional connectomes in the datasets (twice the number of participants). Therefore, by definition, using the number participants as the number of components accounts for 100% of the variance in the data. As a matter of fact, the number of participants correctly identified is close to the best number of PCs. However, in the spinal cord it seems that there are many more similarities across participants, probably due to imaging and anatomical constraints, thus the number of PCs is smaller than the actual number of individuals in the datasets.

By reconstructing the functional connectivity (FC) matrices using the number of Principal Components corresponding to the peak $Idiff$ score, we generated inputs for the fingerprinting pipeline and computed the identifiability matrix for each dataset, as shown in Figure S3. This reconstruction led to a general improvement in both scores and accuracy. For *Dataset 1*, we correctly identified 14 out of 18 participants (1 up from 13 using the raw FC), with a Cohen's d of 2.79. In *Dataset 2*, the number of correctly identified participants increased from 8 to 11 out of 43 (Cohen's d = 1.17). Similarly, for *Dataset 3*, identification improved from 10 to 13 out of 15 participants, with a Cohen's d of 1.71.

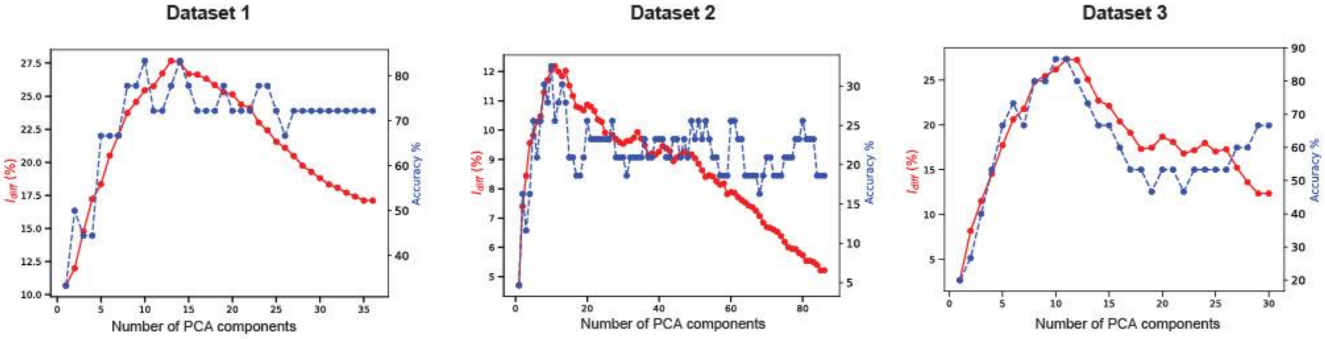


**Figure S2.** $Idiff$ values in percentage (in red) and accuracy of subjects identification with respect to the number of PCA components. The maximum value is close to the number of participants in Dataset 1 and 3, while Dataset 2 (that also performed overall worse) has a very small best number of components (18) with respect to the number of participants (43).

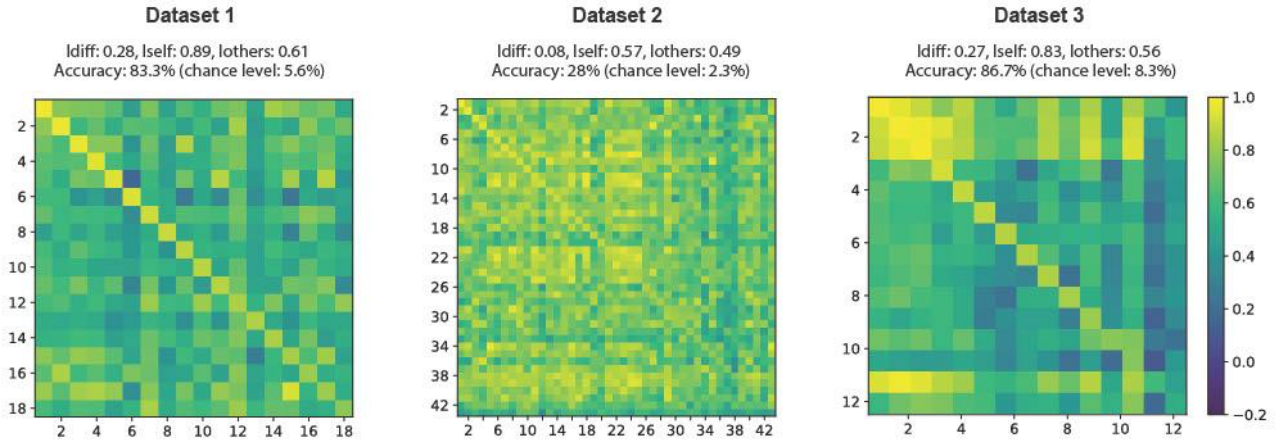


**Figure S3.** Identifiability matrices computed using the optimal number of principal components (PCs) that maximizes $Idiff$.

### Fair brain comparison

#### *HCP dataset*

To ensure a fair comparison, we computed brain identifiability ($Idiff$) scores using data from the HCP dataset, separately from our spinal cord data. Importantly, these analyses were not combined across modalities or datasets, but rather performed in parallel. The goal of this analysis was to assess whether the lower identifiability observed in the spinal cord could be partly explained by differences in data length rather than by intrinsic modality-specific properties. To this end, we matched both the number of time points and individuals used in our spinal cord analyses. We applied the same Schaefer 119 parcellation across datasets. Since the HCP dataset contains 1200 time points, we segmented the data into 7 windows of T time points, with T adjusted to align with the time resolution of each spinal cord dataset. This approach yielded Idiff scores that were comparable to those observed for the spine-print, suggesting that the number of time points significantly influences identifiability performance. This observation aligns with findings from Van De Ville et al. (2021), who systematically demonstrated that brain fingerprint accuracy improves with longer temporal windows.

We report the results of this comparison in Table S1, indicating the mean across the 7 windows of the *Idiff, Iself* and *Iothers* scores as well as the participants identifiability accuracy (PIA) with the number of participants correctly identified.

| | **Spinal cord scores** | **HCP scores** |
|---|---|---|
| **Dataset 1**<br>**T = 288**<br>**Ns = 18** | $Idiff$: **0.17**<br>$Iself$: 0.64<br>$Iothers$: 0.46<br>PIA: **72.2% (13)** | $Idiff$: 0.15<br>$Iself$: 0.48<br>$Iothers$: 0.33<br>PIA: **72.2% (13)** |
| **Dataset 2**<br>**T = 250**<br>**Ns = 43** | $Idiff$: 0.06<br>$Iself$: 0.37<br>$Iothers$: 0.31<br>PIA: 20.9% (9) | $Idiff$: **0.13**<br>$Iself$: 0.46<br>$Iothers$: 0.33<br>PIA: **62.8% (27)** |
| **Dataset 3 (spine)**<br>**T = 230**<br>**Ns = 15** | $Idiff$: 0.09<br>$Iself$: 0.59<br>$Iothers$: 0.50<br>PIA: 53.3% (8) | $Idiff$: **0.16**<br>$Iself$: 0.47<br>$Iothers$: 0.31<br>PIA: **73.3% (11)** |

**Table S1.** Identifiability scores ($Idiff$), self-similarity ($Iself$), between-subject similarity (Iothers), and Participant Identification Accuracy (PIA) for spinal cord and HCP scores across the three datasets.

***Dataset 3***

After having examined the HCP dataset restricted to the brain, we next returned to Dataset 3 (brain + spinal cord) to test whether similar effects would hold when considering functionally relevant systems. Since the spinal cord is predominantly motor-related, we compared it against the somatomotor network (SM) rather than unrelated systems such as the visual network.
Moreover, to ensure a fair comparison based on the functional properties of the two structures, we also calculated fingerprint scores focusing solely on the somatomotor network

(SM) and we obtained an $Idiff$ of 0.18 (Cohen's D: 1.44) obtaining an accuracy of individual identification of 53.3% (8 individuals out of 15). We then combined the SM ROIs with the spinal cord, yielding an $Idiff$ of 0.16 and a very large Cohen's D of 1.87, without any change in the accuracy of individuals identification: 93.3%, 14 out of 15, reaching exactly the same performance as using all brain ROIs. The reader is addressed to Figure S4. In contrast, as expected from literature, using only the visual cortex, we obtain a drop in both $Idiff$ (reaching 0.09) and accuracy: 6 participants were correctly identified (40%).

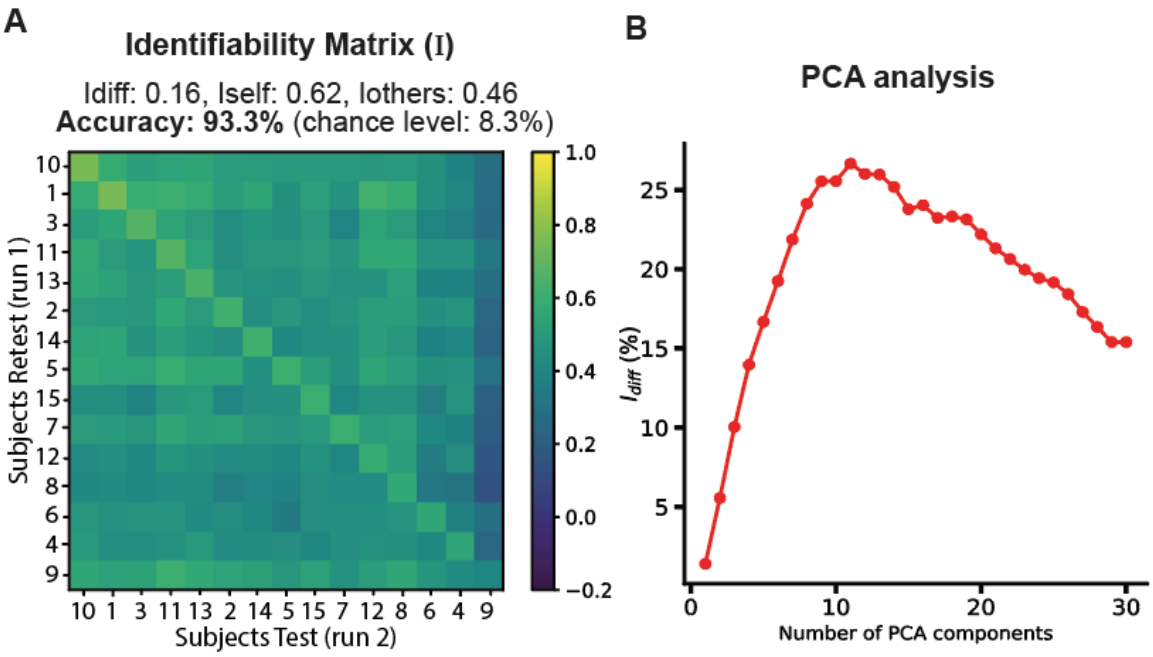


**Figure S4.** (A) Identifiability matrix using the 15 somatomotor bilateral brain regions and the spinal cord regions (N= 98). Scores are reported in the subtitle. (B) PCA analysis of the Idiff scores optimization reaches up to 0.267 (26.7%) for 11 principal components.

**Linear regression method: investigating brain and spinal cord time courses**

Supplementary Figure S5 shows the matricial scheme of the linear regression, displaying all regression elements in matrix form, with the fingerprint input data highlighted in red.
Residuals are calculated by subtracting the predicted values from the original time series: $\epsilon_{sc\backslash br}$ = $Y_{sc}$ - $\widehat{Y_{sc}}$ represents spinal residuals after removing brain-predicted components, while $\epsilon_{br\backslash sc}$ = $Y_{br}$ - $\widehat{Y_{br}}$ represents brain residuals after removing spinal-predicted components.

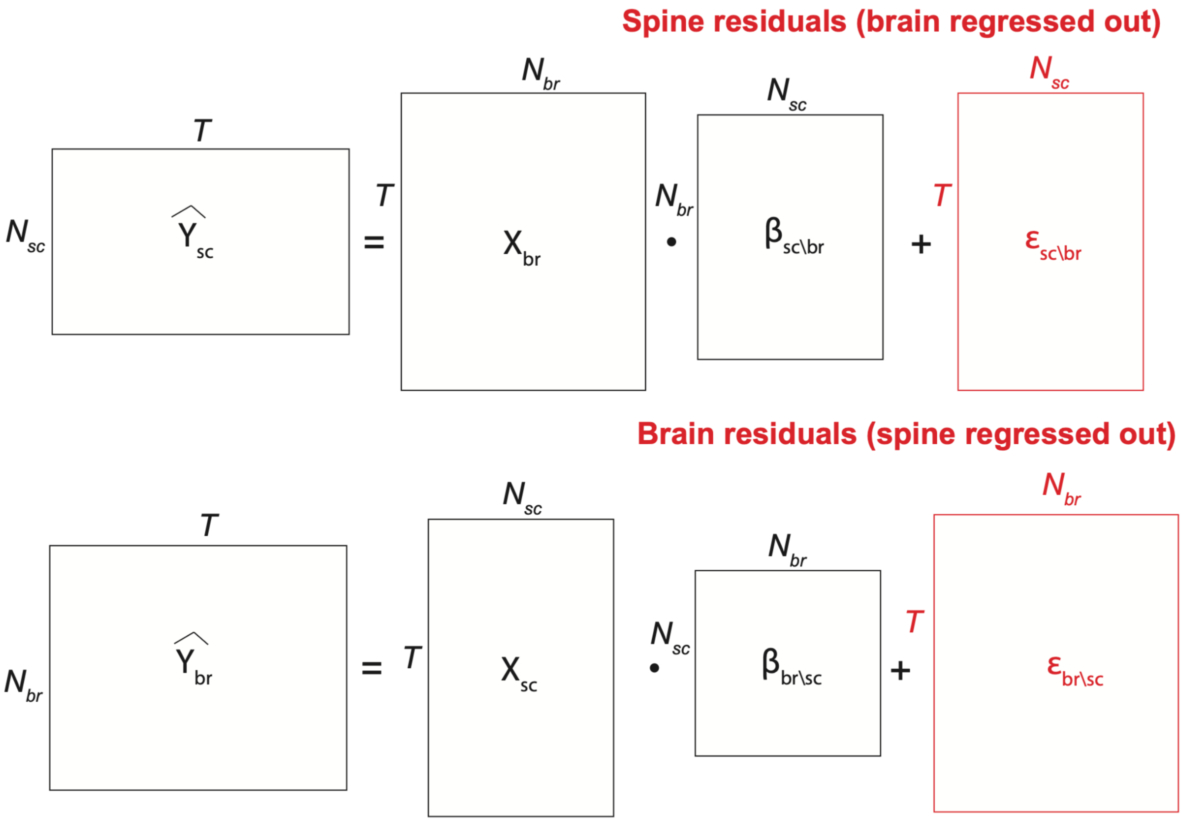


**Figure S5. Linear regression.** Matrix representation showing predicted spinal time courses $\widehat{Y_{sc}}$ as a linear regression of brain time series $X_{br}$, and $\widehat{Y_{br}}$ as a linear predictor of spinal activity $X_{sc}$. The red matrix indicates input data for the fingerprint analysis pipeline.
